# Non-redundant dimethyl sulfoxide reductases influence *Salmonella enterica* serotype Typhimurium anaerobic growth and virulence

**DOI:** 10.1101/2022.11.29.517730

**Authors:** E Cruz, AL Haeberle, TL Westerman, ME Durham, MM Suyemoto, LA Knodler, JR Elfenbein

## Abstract

Facultative anaerobic enteric pathogens can utilize a diverse array of alternate electron acceptors to support anaerobic metabolism and thrive in the hypoxic conditions within the mammalian gut. Dimethyl sulfoxide (DMSO) is produced by methionine catabolism and can act as an alternate electron acceptor to support anaerobic respiration. The DMSO reductase complex consists of three subunits, DmsA, DmsB, and DmsC, and allows bacteria to grow anaerobically with DMSO as an electron acceptor. The genomes of non-typhoidal *S. enterica* encode three putative *dmsABC* operons, but the impact of the apparent genetic redundancy in DMSO reduction on fitness of non-typhoidal *S. enterica* during infection remains unknown. We hypothesized that DMSO reduction would be needed for *S. enterica* serotype Typhimurium (*S*. Typhimurium) to colonize the mammalian gut. We demonstrate a *S.* Typhimurium mutant with loss of function in all three putative DMSO reductases (Δ*dmsA^3^*) poorly colonizes the mammalian intestine when the microbiota are intact and when inflammation is absent. DMSO reduction enhances anaerobic growth through non-redundant contributions of two of the DMSO reductases. Furthermore, DMSO reduction influences the expression of the type-3 secretion systems needed for virulence. Collectively, our data demonstrate that the DMSO reductases of *S.* Typhimurium are functionally non-redundant and suggest DMSO is a physiologically relevant electron acceptor that supports *S. enterica* fitness in the gut.

## Introduction

Non-typhoidal *Salmonella enterica* are a leading cause of bacterial food-borne enterocolitis world-wide (1, 2). In addition to human disease, *Salmonella* can colonize and cause disease in many species of animals. There are more than 2600 *Salmonella* serotypes, which can be divided into functional groups based on the common disease manifestation in infected individuals. Disease manifestations depend on host immune status as well as bacterial characteristics that dictate host and tissue tropism. Host-restricted *Salmonella* serotypes (extraintestinal pathovars) largely cause sepsis in a limited range of hosts while generalist serotypes (gastrointestinal pathovars) cause enterocolitis, characterized by severe neutrophilic inflammation and inflammatory diarrhea in many host species (3). When compared with extraintestinal pathovars, the genomes of gastrointestinal pathovars are enriched with operons dedicated to anaerobic metabolism, demonstrating the importance of anaerobic respiration in enteric salmonellosis (3).

Anaerobic metabolism is critical for enteric pathogens like *Salmonella enterica* to colonize the intestine. Oxygen tension declines along the length of the gut, mediated in large part by oxygen consumption by facultative anaerobes residing in the intestinal lumen (4, 5). Fermentation of complex carbohydrates by anaerobes produces butyrate, which fuels colonocyte oxygen consumption and ensures oxygen tension declines from the epithelial surface to the lumen (6). Because oxygen is limiting in the intestinal lumen, alternate electron acceptors such as nitrate and fumarate are also employed to facilitate the luminal growth of facultative anaerobes (7, 8). However, the inflammatory phase of enteric salmonellosis changes the availability of electron acceptors that support respiration within the gut. Intestinal inflammation increases luminal oxygenation while also providing the anaerobic electron acceptors nitrate and tetrathionate, all of which enhance the luminal replication of *Salmonella* in the mammalian intestine (6, 9–11). Although it is known that tetrathionate and nitrate respiration support *Salmonella* growth in the inflamed intestine, the impact of respiration using the alternate electron acceptor dimethyl sulfoxide (DMSO) in *Salmonella* gut colonization is unknown.

Dimethyl sulfide (DMS) plays an important role in the global sulfur cycle (12). It is found in aquatic and terrestrial environments and in the mammalian gastrointestinal tract (13–21). In the gut, the production of DMS occurs from microbial catabolism of sulfated amino acids that are not absorbed by the host. Catabolism of methionine and cysteine produce methanethiol and hydrogen sulfide which are both toxic to gut mucosa at high concentrations (22). Although the colonic mucosa primarily oxidizes methanethiol and hydrogen sulfide to produce thiosulfate (23), methanethiol and hydrogen sulfide may also be converted into DMS (13). Dimethyl sulfide can then be oxidized into DMSO or dimethylsulfone (DMSO_2_) by both enzymatic and non-enzymatic processes (13). DMS and DMSO can also be found in some fruits, vegetables, grains, and beverages, suggesting a dietary source for both compounds (13, 24). Therefore, it is plausible that DMSO could serve as an alternate anaerobic electron acceptor to support *Salmonella* growth within the intestine.

In this work, we test the central hypothesis that DMSO reduction supports *Salmonella enterica* serovar Typhimurium gut colonization. We demonstrate a *Salmonella* mutant lacking all three annotated putative *dmsA* homologs exhibits decreased gut colonization in bovine and murine infection models, likely due to its role in supporting anaerobic growth. Using *trans*-complementation, we show that two of the three *dmsA* homologs can enhance anaerobic growth in the presence of DMSO and demonstrate they lack functional redundancy *in vitro*. In addition to its role in supporting anaerobic growth, we document a role for DMSO and its reduced form, DMS, in titration of virulence gene expression. Taken together, our work demonstrates DMSO reduction is a part of the metabolic repertoire used by *Salmonella* for pathogenesis in the mammalian gut.

## Results

### DMSO reduction supports colonization of the anaerobic gut

DMSO reductases support anaerobic respiration using DMSO as a terminal electron acceptor (25). The DMSO reductase is a trimeric protein complex composed of the periplasmic DmsA, the catalytic subunit that binds and reduces DMSO to DMS, the periplasmic electron transfer subunit DmsB, and the cytoplasmic membrane anchor subunit DmsC that binds and oxidizes the electron donor, menaquinol (26). The *Salmonella* Typhimurium genome encodes three putative *dmsABC* operons (**Figure 1A**): *STM0964-STM0966* (*STM14_1089-1091*), *STM2530-2527* (*STM14_3103-3100*), and *STM4305-STM4308 (STM14_5178-5181)* (*3*). The DMSO reductase encoded by *STM0964*-*0966* has comparable genomic context (**Figure 1A**) and high amino acid identity (90-94%) with *dmsABC* in the close relative *E. coli* (**Figure 1B**), suggesting *STM0964-STM0966* is likely to be an ancestral copy of *dmsABC*. *STM2530-STM2527* and *STM4305-STM4308* each encode a fourth gene within their predicted operons (**Figure 1A**) and share lower amino acid identity with *E. coli dmsABC* (**Figure 1B**), suggesting these operons may have been acquired after *Salmonella* and *Escherichia* diverged from their common ancestor. Both gastrointestinal and extraintestinal *Salmonella* serotypes encode intact copies of an operon homologous to *STM0964-STM0966*, but some extraintestinal serotypes have coding sequence disruptions within the other two operons (3), leading us to hypothesize that DMSO reduction is important for *Salmonella* Typhimurium gastrointestinal colonization.

**Figure 1:**
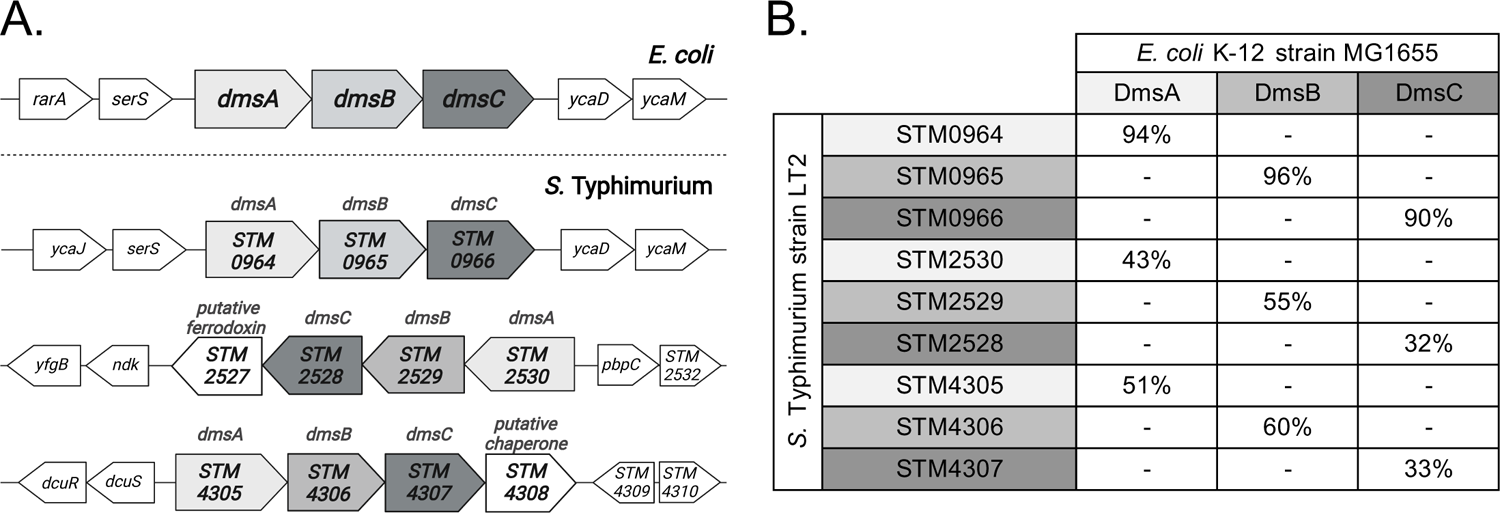
The *Salmonella* Typhimurium genome encodes three operons homologous to *dmsABC* of *E. coli*. (A) Organization and genetic context of the three putative DMSO reductase operons in *S.* Typhimurium strain LT2 and *dmsABC* in *E. coli* K-12 strain MG1655. Genomic contexts were adapted from Biocyc (65). (B) Percent amino acid identity shared between each DmsABC homolog in *S.* Typhimurium strain LT2 and *E. coli* K-12 strain MG1655. Amino acid identities were calculated from sequence alignments generated with Clustal Omega (66). The figure was created using Biorender.com.

We utilized the calf ligated ileal loop model to determine the requirement for DMSO reduction during enteric salmonellosis. In the calf model, *S.* Typhimurium infection causes acute, severe neutrophilic enteritis which closely resembles human disease (27, 28). Induction of severe neutrophilic inflammation and luminal fluid accumulation requires the action of five effector proteins secreted by the type-3 secretion system-1 (T3SS1) (29). We hypothesized that DMSO reduction would support *S.* Typhimurium intestinal colonization in the presence of neutrophilic inflammation, but not in its absence. Intestinal segments were inoculated with an equal mixture of the WT and a mutant lacking all three annotated *dmsA* homologs (Δ*dmsA^3^*; Δ*STM0964*Δ*STM2530*Δ*STM4305*) or with an equal mixture of a mutant lacking *Salmonella* Pathogenicity Island-1 (ΔSPI1), which encodes the T3SS1, and a ΔSPI-1Δ*dmsA^3^* mutant as an inflammation-deficient condition. We observed a significant colonization defect for the Δ*dmsA^3^* mutant in small intestinal tissue in the ΔSPI-1 genetic background but not in the WT genetic background (**Figure 2A**). Significant luminal fluid accumulated only in loops inoculated with the WT organism, but not in loops inoculated with organisms in the ΔSPI-1 background (**Figure 2B**). Fluid accumulation correlates with neutrophilic inflammation in this model (29), so the lack of fluid accumulation in loops inoculated with organisms in the ΔSPI-1 genetic background confirms no substantial inflammation was present in these loops. These data led us to reject our hypothesis and suggested that DMSO reduction is important for tissue colonization in the absence of inflammation.

**Figure 2:**
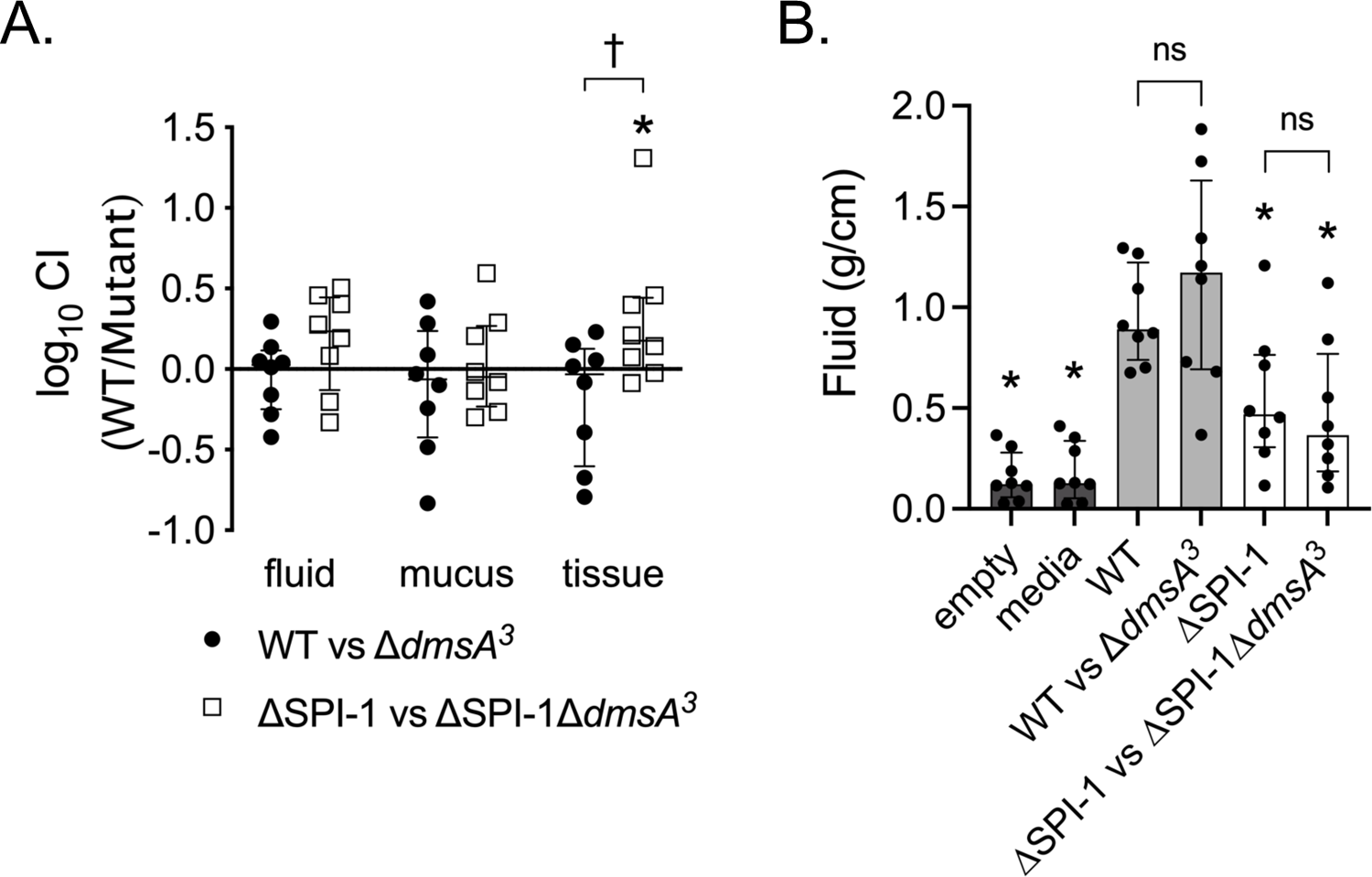
Deletion of all three *dmsA* homologs impairs colonization of bovine ileal tissue in the absence of inflammation. Ligated intestinal loops were inoculated with ∼10^9^ CFU of an equal mixture of the WT and Δ*dmsA*^3^ mutant in inflammatory (WT) or non-inflammatory (ΔSPI-1) genetic backgrounds. At 12 hours post-infection, calves were euthanized, and intestinal segments were harvested. (A) Luminal fluid, mucus, and intestinal tissue were separated, homogenized, diluted, and plated to determine competitive index (CI). Each symbol represents one loop. * denotes significant difference in CI (p<0.05) while † denotes significant difference between genetic backgrounds (p<0.05). (B) Accumulated luminal fluid was weighed and normalized to the starting length of each loop. * indicates significant difference in fluid accumulation compared with the WT-infected loop (p<0.05), ns indicates no difference. Individual points represent a single animal with median +/- interquartile range indicated. Statistical differences were detected by repeated-measures ANOVA with Tukey’s (A) or Dunnett’s (B) correction for multiple comparisons.

Next, we hypothesized that an intact microbiota would be needed for DMSO reduction to support *Salmonella* gut colonization. We used two different murine salmonellosis models to test our hypothesis. First, we used an oral infection model in which *Salmonella*-resistant mice shed organisms in feces for more than 40 days (30). Importantly, the microbiota are not disturbed prior to *Salmonella* infection in this model and mice develop mild tissue inflammation after 7 days’ infection (31). We observed a competitive disadvantage for the Δ*dmsA^3^* mutant in the cecum of orally infected mice (**Figure 3A**). Next, we used the murine colitis model in which *Salmonella*-susceptible mice are treated with a high dose of streptomycin to disrupt the microbiota which allows robust *Salmonella* replication in the cecum and the subsequent development of severe neutrophilic inflammation (32). We observed no colonization defect for the Δ*dmsA^3^* mutant in either the inflammatory (WT) or non-inflammatory (ΔSPI-1) genetic backgrounds after antimicrobial disruption of the gut microbiota (**Figure 3B**). Altogether, our data suggest that DMSO reduction supports intestinal tissue colonization in the mammalian gut when the microbiota are intact.

**Figure 3:**
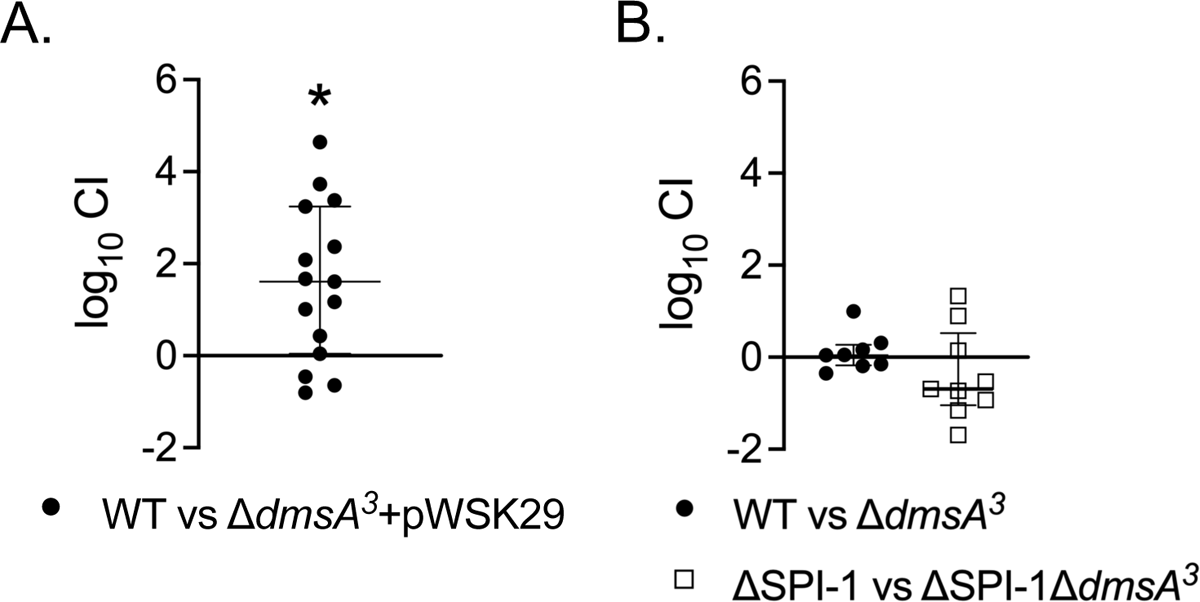
Deletion of all three *dmsA* homologs impairs colonization of the murine cecum with an intact microbiota. (A) CBA/J mice were inoculated with ∼10^8^ CFU of an equal mixture of WT and Δ*dmsA^3^ +* pWSK29 mutant by gavage and euthanized 8-days post-infection. Data are combined from two separate experiments. (B) C57BL/6 mice were treated with 20 mg of streptomycin by gavage and then inoculated with ∼10^8^ CFU of an equal mixture of WT and Δ*dmsA^3^* or ΔSPI-1 and ΔSPI-1Δ*dmsA^3^* one-day later. Mice were euthanized 4-days post-infection. The cecum was harvested from each mouse, homogenized, serially diluted, and plated to determine CI. Data points represent individual animals and the median +/- interquartile range is indicated. * denotes significant difference in CI as determined by Student’s t-test (p<0.05).

Since DMSO can act as an electron acceptor to support anaerobic respiration (25), we hypothesized that DMSO reduction would support gut colonization by enhancing anaerobic growth. We grew the WT and the Δ*dmsA^3^* mutant in anaerobic conditions in rich media, with and without DMSO. In the absence of DMSO, both the WT and the Δ*dmsA^3^* mutant grew with similar kinetics (**Figure 4A**). In the presence of DMSO, we observed a significant growth enhancement for the WT organism but not the Δ*dmsA^3^* mutant (**Figure 4B**). Since DMSO could serve as a source of both carbon and sulfur to enhance growth, we assessed the effects of addition of DMS, the end metabolite, to the media. DMS did not alter the growth of either the WT or the Δ*dmsA^3^* mutant, (**Figure 4C**) suggesting that reduction of DMSO was responsible for the observed anerobic growth enhancement. No further growth augmentation was achieved in rich media with the addition of up to 10-fold higher concentrations of DMSO (**Figure 4D**). These data demonstrate that DMSO reduction by one or more of the putative DMSO reductases enhances *Salmonella* anaerobic growth in rich media.

**Figure 4:**
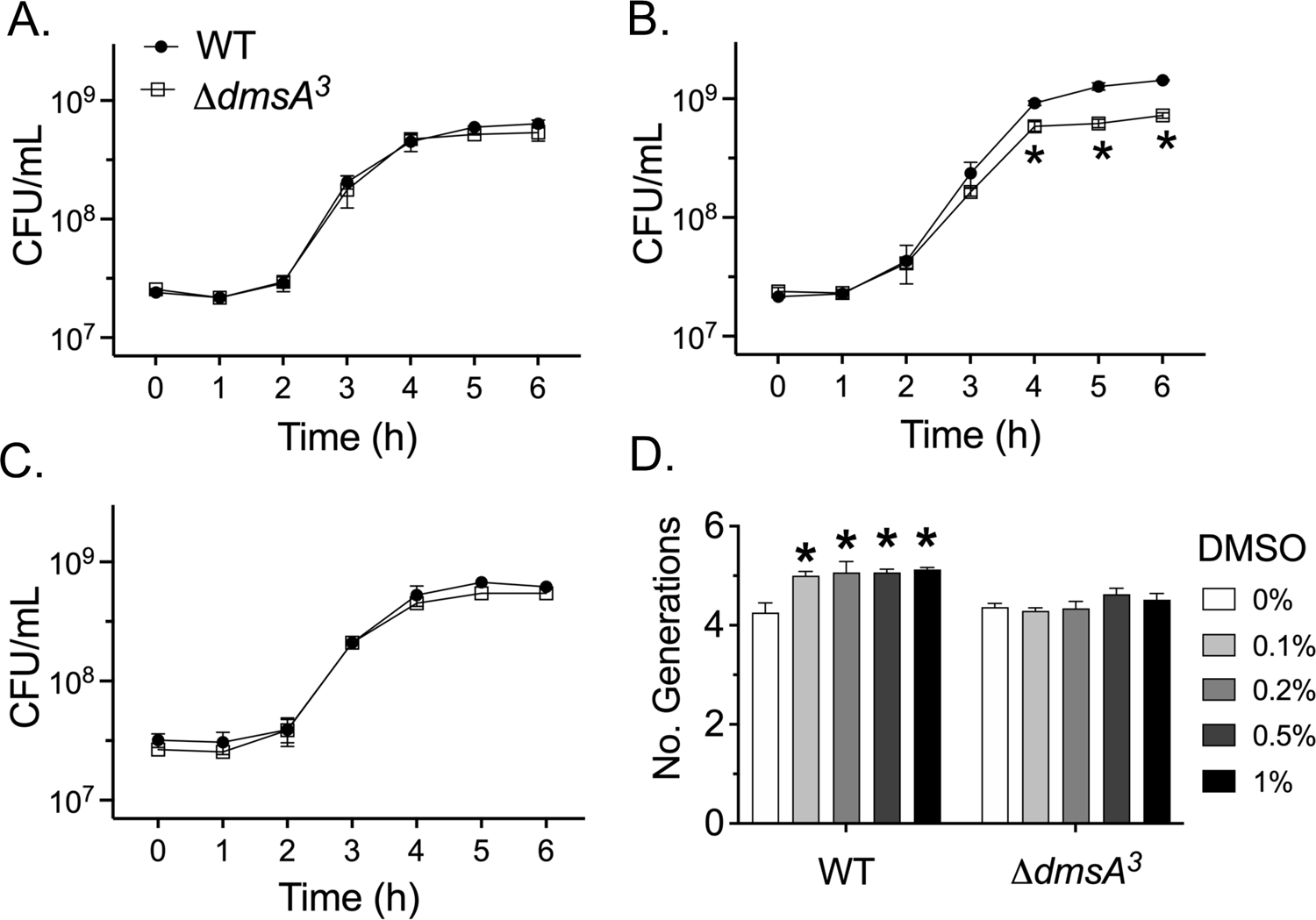
DMSO enhances the anaerobic growth of *Salmonella* Typhimurium in rich media. Overnight cultures were moved into an anaerobic chamber and diluted into (A) LB broth, (B) LB supplemented with 0.1% (v/v) DMSO, (C) LB supplemented with 0.1% (v/v) DMS, or (D) LB supplemented with the indicated concentrations of DMSO (v/v). Bacteria were grown anaerobically at 37°C with aliquots taken at the specified times (A-C) or at 4 hours growth (D). Aliquots were serially diluted and plated to quantify CFU/mL. The number of generations (D) was calculated from CFU/mL measurements. Data points or bars represent the mean +/- SEM of 3 (A-C) and 5 (D) independent experiments. * denotes significant difference (p<0.05) between the WT and mutant as detected by Student’s t-test with Holm-Šídák correction for multiple comparisons (B) or significant difference between DMSO-supplemented LB and LB alone as detected by two-way ANOVA with Dunnett’s correction for multiple corrections (D).

Although DMSO is an electron acceptor that supports anaerobic growth, it is not the preferred anaerobic electron acceptor by the close relative *E. coli* (33). We observed DMSO reduction supported *S*. Typhimurium colonization of bovine intestinal tissue only in the absence of neutrophilic inflammation (**Figure 2A**). One potential explanation for our observation is that preferred electron acceptors such as oxygen, nitrate, and tetrathionate become abundant in the inflamed gut, negating the role of DMSO reduction in supporting *S*. Typhimurium gut colonization (6, 9, 11). To establish whether preferred electron acceptors impact the use of DMSO reduction to enhance anaerobic growth, we performed competitive growth assays in media with DMSO and added oxygen, tetrathionate, or nitrate. Consistent with the role of oxygen as the preferred electron acceptor, the Δ*dmsA^3^* mutant had a significant disadvantage when grown in rich media with DMSO only when oxygen was absent (**Figure 5A**). The Δ*dmsA^3^* mutant growth disadvantage persisted in anaerobic mucin broth supplemented with DMSO, demonstrating the enhanced growth was not media specific (**Figure 5B and 5C**). If either nitrate or tetrathionate were added to anaerobic mucin broth with DMSO, growth of the Δ*dmsA^3^* mutant and WT bacteria were comparable (**Figure 5B and 5C**) but WT bacteria outcompeted the mutants unable to reduce tetrathionate (Δ*ttrA)* and nitrate (Δ*napA*Δ*narZ*Δ*narG*), as expected. These data suggest DMSO reduction enhances *Salmonella* growth only in the absence of oxygen, tetrathionate, and nitrate, which are preferred electron acceptors that are available during *Salmonella*-induced intestinal inflammation.

**Figure 5:**
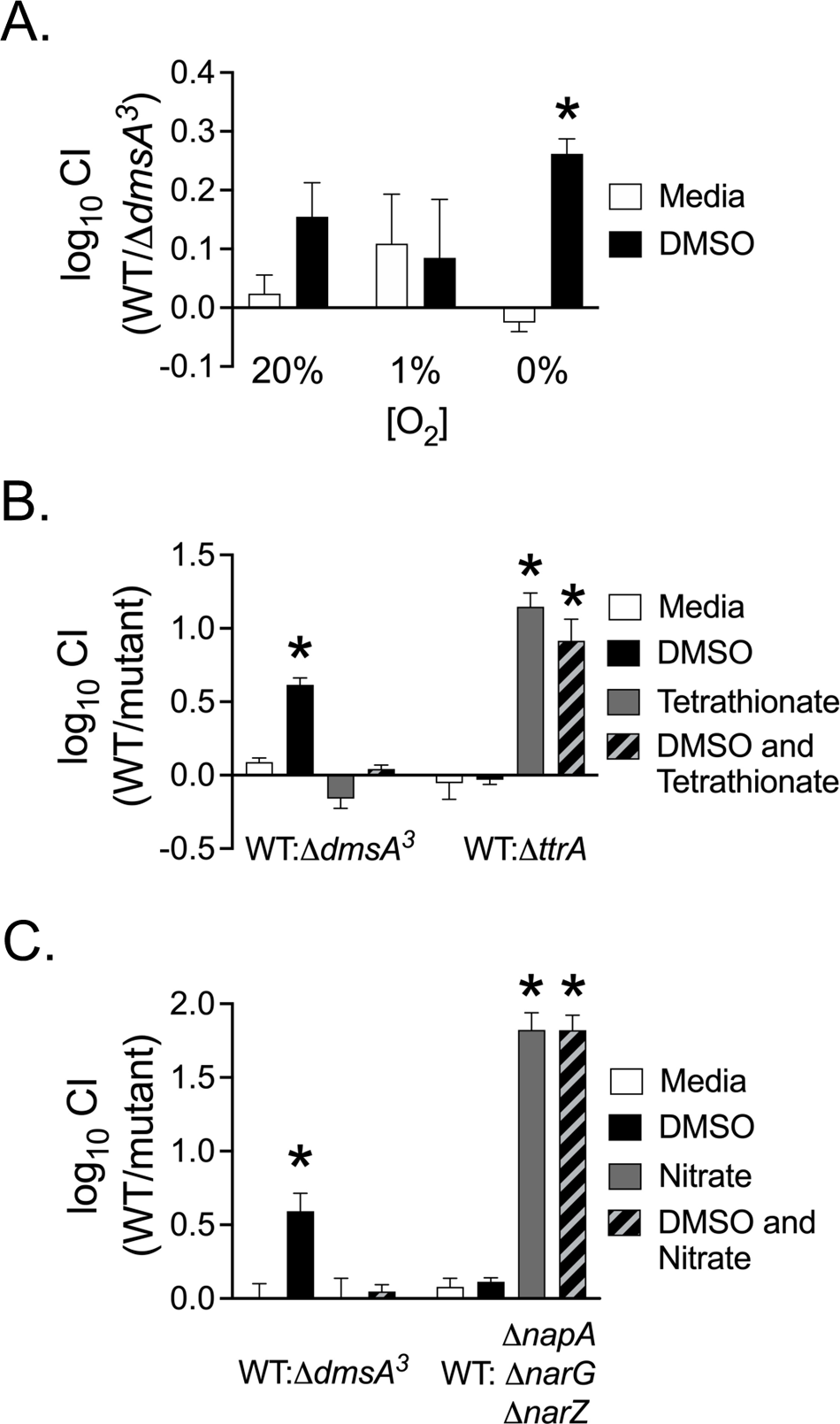
The growth advantage provided by DMSO is lost when oxygen, tetrathionate, or nitrate is available. (A) An equal mixture of the WT and Δ*dmsA*^3^ mutant was diluted into LB +/- 1% (v/v) DMSO and incubated standing at 37°C with the indicated oxygen tensions for 6 hours. (B-C) Equal mixtures of the indicated bacterial strains were diluted into pre-reduced 0.5% mucin broth supplemented with 40 mM of the indicated electron acceptor and incubated in an anaerobic chamber at 37°C for 18 hours. CI was determined at 6 hours (A) or 18 hours (B-C) growth. Bars represent the mean +/- SEM of 5 (A), 3 (B), or 4 (C) independent experiments. * denotes significant difference in CI by Student’s t-test (p<0.05).

### The three *dmsA* homologs are functionally non-redundant

Next, we sought to establish the role of each DMSO reductase individually to support *S*. Typhimurium anaerobic growth. First, we tested the impact of each individual annotated DMSO reductase on the response of the Δ*dmsA^3^* mutant to DMSO through complementation *in trans*. Complementation *in trans* with either *STM0964* or *STM4305* restored the Δ*dmsA^3^* mutant growth to WT levels in the presence of DMSO (**Figure 6A**). However, no DMSO-dependent growth enhancement was observed when *STM2530* was added *in trans* to the Δ*dmsA^3^* mutant (**Figure 6A**). Next, we sought to confirm our complementation studies by testing the anaerobic growth of targeted mutants lacking one or more *dmsA* homologs. A double mutant in Δ*STM0964ΔSTM4305* was selected first for study because *STM0964 and STM4305* both reversed the phenotype of the Δ*dmsA^3^* mutant. As we observed for the *ΔdmsA^3^* mutant, the *ΔSTM0964ΔSTM4305* mutant did not gain a growth advantage in the presence of DMSO (**Figure 6B**). Treatment with DMSO did not enhance the anaerobic growth of the Δ*STM0964* mutant, but the Δ*STM4305* mutant exhibited a DMSO-dependent growth enhancement similar to that observed for WT bacteria (**Figure 6B**). The Δ*STM0964* mutant complemented *in trans* with *STM0964* restored the ability to respond to DMSO at WT levels, as expected (**Figure 6B**). However, complementation *in trans* with *STM4305* did not restore a growth enhancement to the Δ*STM0964* mutant in the presence of DMSO (**Figure 6B**). These data suggest that *STM0964* is the dominant DMSO reductase for anaerobic DMSO respiration and that *STM4305* can act as an alternate DMSO reductase to enhance anaerobic growth with DMSO, when it is the sole DMSO reductase. However, *STM4305* cannot functionally replace *STM0964*.

**Figure 6:**
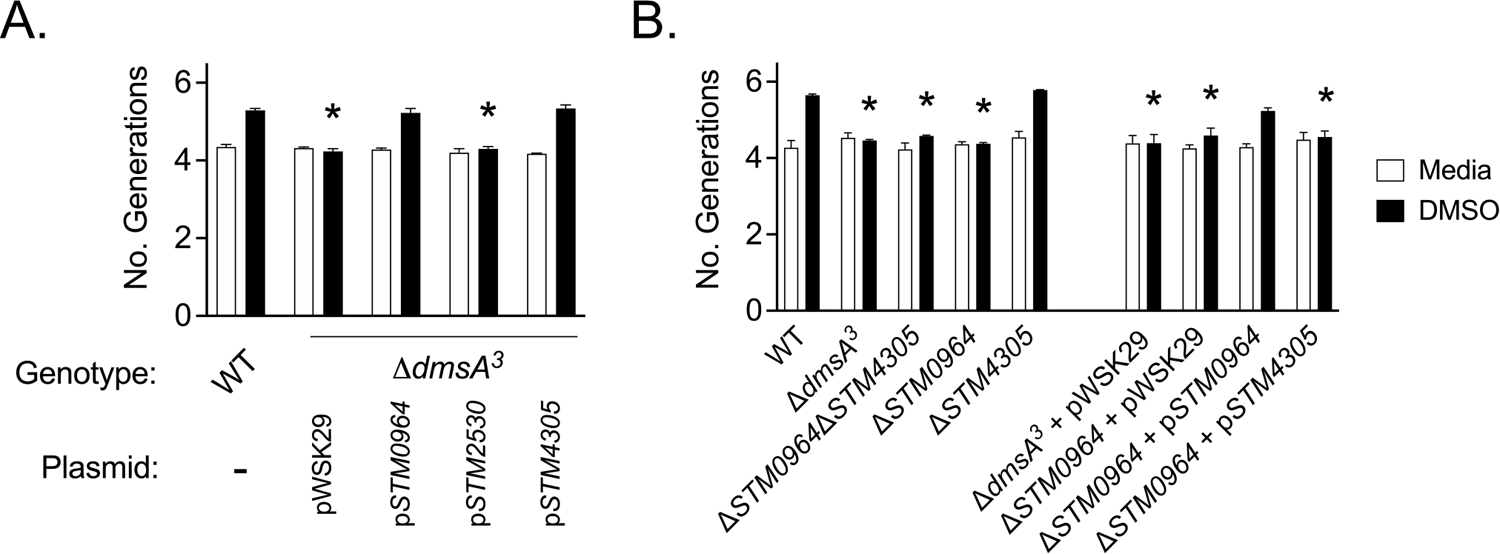
*STM0964* and *STM4305* have non-redundant functions in anaerobic DMSO respiration. Overnight cultures were moved into an anaerobic chamber and diluted into pre-reduced LB broth +/- 0.1% (v/v) DMSO. The number of generations was determined after 4 hours growth at 37°C. Bars represent the mean +/- SEM of 3 independent experiments. * denotes significant difference between the WT and indicated strain within a given condition, as determined by two-way ANOVA with Dunnett’s correction for multiple comparisons (p<0.05).

### DMSO reduction influences virulence gene expression

Previous work demonstrated that DMS, the product of DMSO reduction, inhibits *S*. Typhimurium expression of T3SS1 genes and subsequent invasion into tissue-cultured epithelial cells (34). However, no link was made between DMSO reduction and suppression of T3SS1 activity. Based on the reported observations, we hypothesized that DMSO would have no impact on T3SS1 gene expression and epithelial invasion in a Δ*dmsA^3^* mutant. We used reporter plasmids with the promoter for *prgH*, a component of the T3SS1 apparatus, driving the expression of *lacZY* to measure the effect of DMSO and DMS on T3SS1 expression in aerobic exponential growth *in vitro*. We observed that addition of both DMSO and DMS significantly reduced activity of the *prgH* promoter for both the WT and the Δ*dmsA^3^* mutant (**Figure 7A**). Consistent with prior reports (34), DMS was a more potent suppressor of T3SS1 promoter activity than DMSO. DMSO treatment also reduced invasion into intestinal epithelial cell (HCT116) monolayers for both the WT and Δ*dmsA^3^* mutant (**Figure 7B**), confirming that reduced *prgH* promoter activity corresponded to reduced T3SS1 function, thereby impacting bacterial invasion into this non-phagocytic cell line. Since expression of T3SS1 genes is downregulated 1-2 hours after bacterial internalization into non-phagocytic cells (35), we tested whether DMSO treatment impacts intracellular T3SS1 expression in mammalian cells. We observed no impact of DMSO treatment on the kinetics of T3SS1 downregulation in HCT116 epithelial cells for either strain (**Figure 7C**).

**Figure 7:**
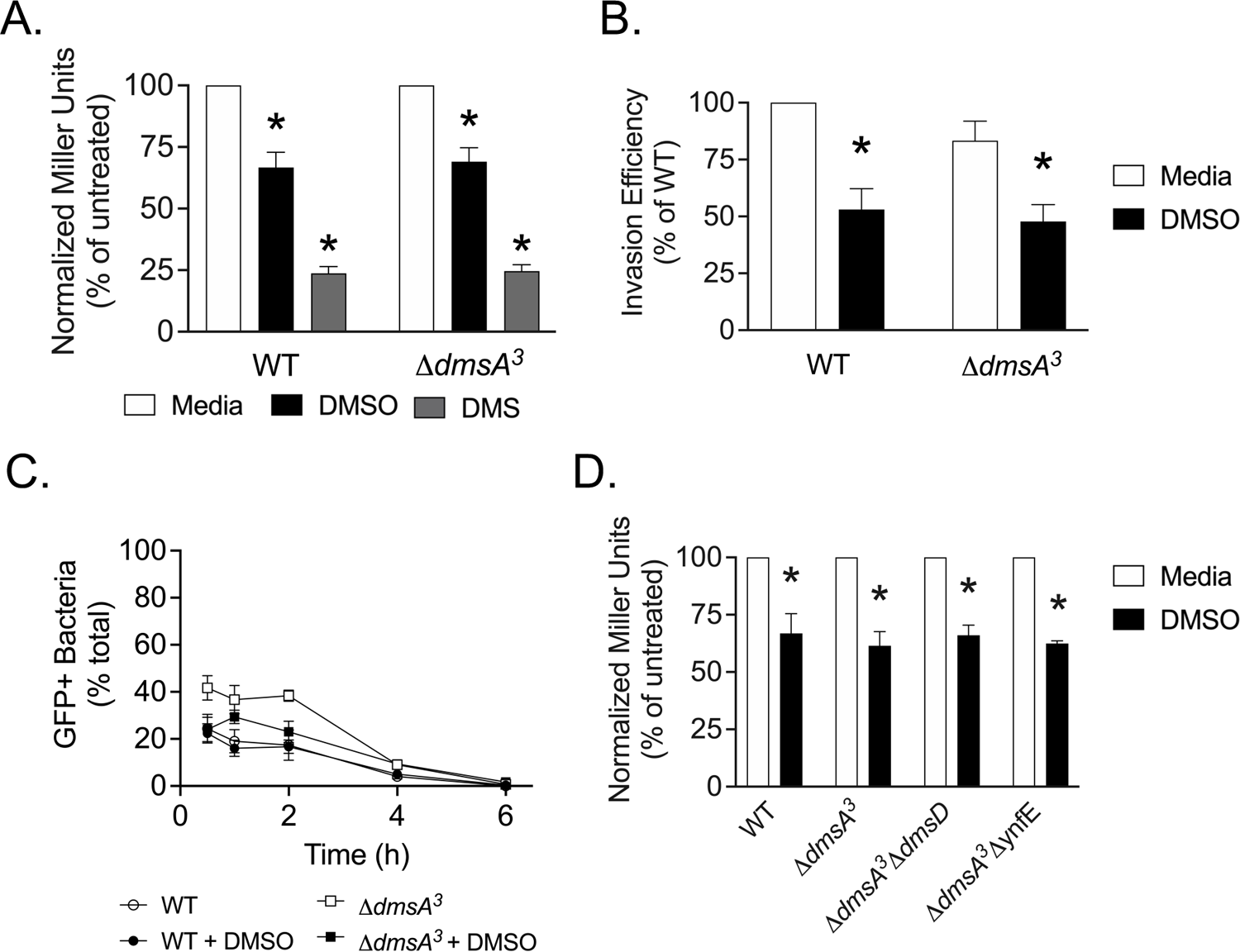
DMSO decreases *prgH* promoter activity and invasion efficiency. (A, D) Overnight cultures of the indicated bacterial strains bearing a P*_prgH_*-*lacZY* transcriptional fusion were diluted into LB and grown aerobically at 37°C. Where indicated, LB was supplemented with 1% (v/v) DMSO or DMS after 1.5 hours growth and incubation continued for another 1.5 hours. The activity of the *prgH* promoter was estimated by measuring β-galactosidase activity, which was converted into Miller Units. (B) Bacteria were prepared as in (A) and invasion efficiency into HCT116 epithelial cell monolayers was determined by gentamicin protection assay at 1 hour post-infection. Invasion efficiency was normalized to the untreated WT strain. (B) HCT116 cells were infected with the indicated bacterial strains (constitutively expressing *mCherry*) bearing a P*_prgH_*-GFP[LVA] transcriptional fusion. Cells were fixed at the indicated timepoints, and the proportion of GFP-positive bacteria quantified by fluorescence microscopy. Bars represent the mean +/- SEM of 3 (A, C, D) or 4 (B) independent experiments. Asterisks (*) indicate significant difference from media alone as detected by Student’s t-test (p<0.05).

Our unexpected observation that DMSO significantly reduced T3SS1 activity in the Δ*dmsA^3^* mutant suggested that there may be another enzyme that can reduce DMSO to produce DMS in aerobic growth, which was how bacteria were grown to induce T3SS1 expression. We made two mutations in the Δ*dmsA^3^* mutant background to test our hypothesis. We deleted the molecular chaperone *dmsD (STM1495* or *STM14_1806)*, which is necessary for DMSO reductase assembly and function (36). We also deleted the *dmsA* paralog *ynfE* (*STM1499* or *STM14_1810*) because the *ynfEFGHI* operon can functionally replace *dmsABC* in *E. coli,* although it has been shown to mediate selenate reduction in *S*. Typhimurium (37, 38). We found decreased activity of the *prgH* promoter in both the Δ*dmsA^3^ΔdmsD* and Δ*dmsA^3^*Δ*ynfE* mutants exposed to DMSO (**Figure 7D**). The observed decrease in *prgH* promoter activity in both mutants was similar to that of the WT and Δ*dmsA^3^* mutant. These data suggest that there is no additional DmsD-dependent DMSO reductase to reduce DMSO in the conditions tested.

The T3SS2 is critical for intracellular survival, particularly in phagocytic cells, and is essential for *S*. Typhimurium virulence (39–41). T3SS2-expressing bacteria are observed in the gut epithelium and phagocytes in the lamina propria by 8 hours infection in the calf model (42). Since we observed a defect for the Δ*dmsA^3^* mutant in intestinal tissue after 12 hours infection in the absence of the T3SS1, we hypothesized that DMSO reduction may enhance intracellular survival. First, we established the impact of DMSO reduction on the activity of a T3SS2 apparatus promoter (P*_ssaG_*). Using *in vitro* conditions designed to mimic those in the lumen of the *Salmonella*-containing vacuole (43), we found DMSO treatment caused an increase in T3SS2 promoter activity for WT bacteria but not the Δ*dmsA^3^* mutant (**Figure 8A**). Similar results were seen upon addition of DMSO to anaerobic rich media (**Figure 8B**), a growth condition that normally represses T3SS2 expression (44). These data suggest that DMSO reduction enhances T3SS2 expression *in vitro*. However, this phenotype was not observed upon infection of J774.1 mouse macrophage-like cells (**Figure 8D**), which would explain why no significant differences in intracellular replication were observed between the WT and Δ*dmsA^3^* mutant either with or without DMSO treatment (**Figure 8C**). Together, these data demonstrate that DMSO reduction increases T3SS2 expression *in vitro*, an effect that has no measurable impact on intracellular bacterial replication *in vitro*.

**Figure 8:**
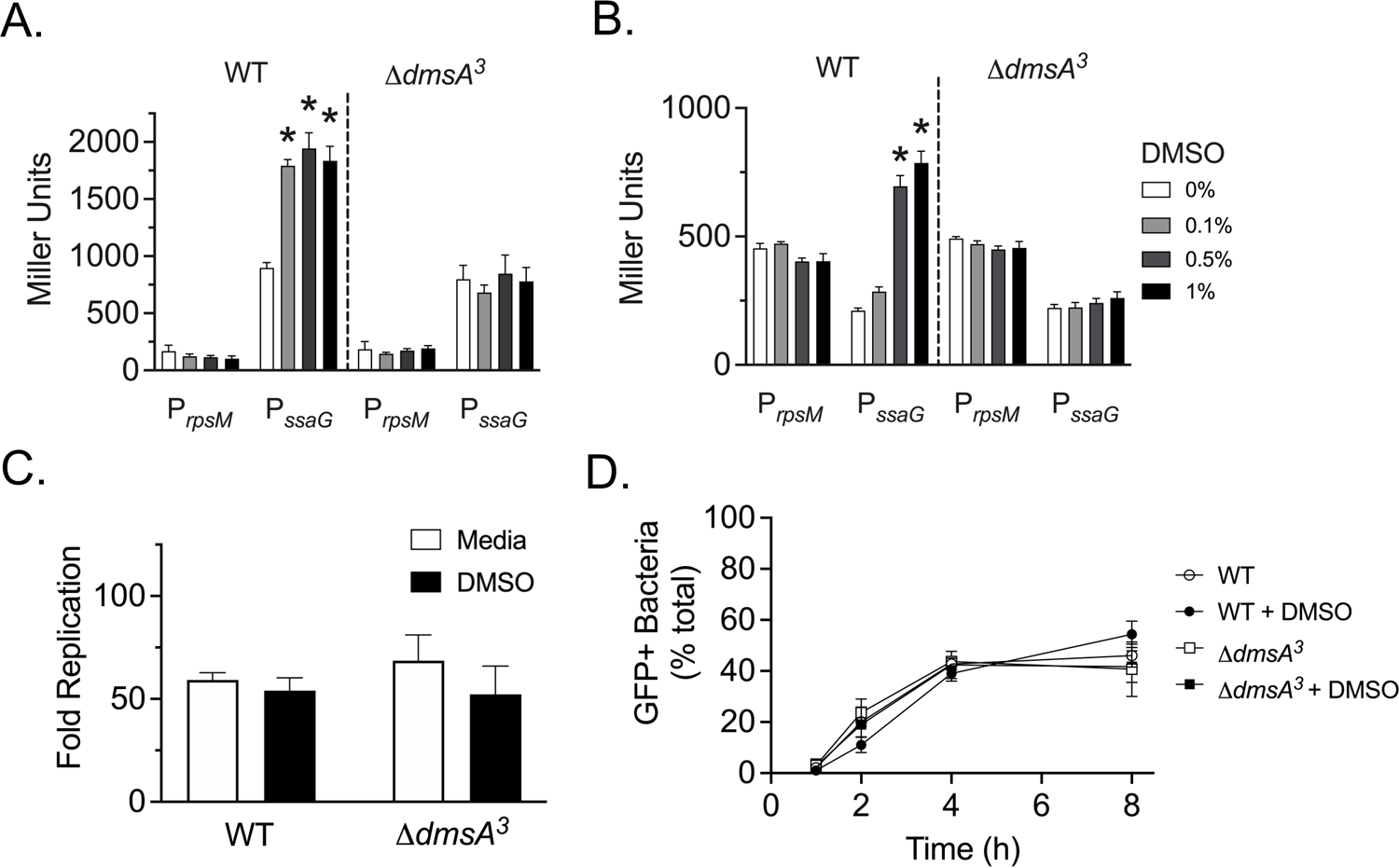
DMSO increases the promoter activity of *ssaG.* (A, B) Overnight cultures of the indicated bacterial strains bearing P*_ssaG_*-*lacZY* transcriptional fusion were diluted into T3SS2-inducing media (A) or into anaerobic pre-reduced LB (B) with the indicated concentrations of DMSO. The activity of the *ssaG* promoter was determined after 24 hours growth. (C) J774A.1 cells were infected with the indicated strains. For DMSO treatment, 1% (v/v) DMSO was added to subcultures for 1.5 hours and at 10 minutes post-infection to J774A.1 cells. Intracellular CFU were quantified by gentamicin protection assay at 1 hour and 16 hours post-infection. Fold-replication was calculated by dividing CFU at 16 hours by CFU at 1 hour. (D) J774A.1 cells were infected with the indicated strains (constitutively expressing *mCherry*) bearing a P*_ssaG_*-GFP[LVA] transcriptional fusion. DMSO was added to subcultures and at 10 minutes post-infection as described in (C). Infected monolayers were fixed at 1, 2, 4, and 8 hours post-infection and the proportion of GFP-positive bacteria was quantified by fluorescence microscopy. Bars represent mean +/- SEM from 3 (A, B, D) or 5 (C) independent experiments. * denotes significant difference as compared with media alone as determined by two-way ANOVA with Dunnett’s correction for multiple comparisons (p<0.05).

## Discussion

In this study, we investigated the role of DMSO reduction in *S.* Typhimurium fitness by characterizing a mutant lacking all three putative *dmsA* homologs. In bovine and murine infection models, disruption of DMSO reduction resulted in decreased colonization of the gastrointestinal tract when the microbiota were intact and in the absence of severe neutrophilic inflammation. DMSO reduction enhanced *Salmonella in vitro* anaerobic growth through the actions of *STM0964* and *STM4305*. Further, we demonstrated DMSO and its reduced form, DMS, both impact expression of virulence paradigms *in vitro*. Together these data support our hypothesis that DMSO reduction impacts pathogenesis by supporting anaerobic metabolism and/or alteration to virulence paradigms during intestinal colonization.

Utilization of the electron acceptor with the highest reduction potential is necessary to maximize ATP synthesis and growth. Oxygen is the preferred electron acceptor in *E. coli*, followed by nitrate and others (33). We observed that DMSO reduction does not enhance *Salmonella* growth *in vitro* when either oxygen, nitrate, or tetrathionate is available in addition to DMSO. One potential explanation for our observation is that growth is maximized when electron acceptors with high redox potentials are available, so no further growth advantage is gained by adding an additional electron acceptor. Another potential explanation is that higher redox potential electron acceptors repress expression of DMSO reductases, as has been demonstrated for oxygen and nitrate in *E. coli* K-12 (45). Our work suggests that tetrathionate is also a preferred electron acceptor over DMSO for *Salmonella*, although future work is needed to establish whether the preference is due to energy generation, regulation, or both.

We demonstrate both *STM0964* and *STM4305* can enhance anaerobic growth in the presence of DMSO. We observed an absolute requirement for *STM0964* to enhance anaerobic growth in the presence of DMSO *in vitro* by complementation *in-trans* of the Δ*dmsA^3^* mutant with *STM0964* and by evaluation of a single deletion mutant lacking *STM0964* alone. These data suggest *STM0964-STM0966* encodes the dominant DMSO reductase that is necessary and sufficient to support anaerobic DMSO reduction. In contrast, the role of the alternate DMSO reductase encoded by *STM4305-STM4307* in enhancing anaerobic growth in the presence of DMSO is less clear. Through complementation of the Δ*dmsA^3^* mutant *in trans* with *STM4305*, we observed an anaerobic growth enhancement in the presence of DMSO, suggesting STM4305 can reduce DMSO to enhance anaerobic growth. However, *STM4305* was unable to *trans*-complement the Δ*STM0964* mutant and deletion of only *STM4305* had no impact on DMSO-dependent anaerobic growth enhancement. These data suggest that *STM4305* is sufficient to enhance anaerobic growth with DMSO when it is the only *dmsA* copy present (or expressed). The apparent discrepancy in the importance of *STM4305* could be explained by increased expression of *STM4305* on the low-copy number plasmid in the complemented Δ*dmsA^3^* mutant (46). However, the lack of phenotype reversal of the Δ*STM0964* mutant by complementation *in trans* with *STM4305* suggests that increased copy number of *STM4305* alone is insufficient for DMSO-dependent anaerobic growth enhancement. We hypothesize that the *dmsA* subunit may be able to associate only with the cognate *dmsBC* subunits, therefore overexpression of only *STM4305* may not generate a functional DMSO reductase complex unless *dmsBC* (*STM4306-STM4307*) are overexpressed at the same time. Taken together, our data demonstrate that the DMSO reductases of *Salmonella* do not exhibit functional redundancy in anoxic conditions *in vitro*.

We were unable to identify a role for *STM2530* in DMSO-dependent anaerobic growth enhancement in rich media. There are several potential reasons for our observations. First, STM2530 shares the lowest amino acid identity with *E. coli* DmsA (43% amino acid identity) of the three annotated DMSO reductases, so it may have higher affinity for a substrate other than DMSO (47). Second, it is possible that STM2530 may reduce DMSO but does not generate sufficient energy for enhanced growth in the conditions tested. Finally, it is possible that *STM2530* is not expressed in the anaerobic rich media we tested, even though we added DMSO to the media. The three nitrate reductases are differentially regulated by oxygen and nitrate concentrations in both *E. coli* and *Salmonella* (9, 48, 49), lending support to the hypothesis that regulation of *STM2530* expression may differ from the other *dmsA* homologs. The genomes of the extraintestinal serotypes *S.* Typhi and *S.* Paratyphi A have frameshift mutations in their *STM2530* homologs (3), suggesting *STM2530* may facilitate intestinal infection or persistence in non-mammalian hosts. Therefore, further work is needed to determine the preferred substrate for STM2530 and conditions activating the expression of *STM2530*.

*Salmonella* has evolved complex regulatory circuits that sense environmental stimuli to titrate the expression of both T3SS. T3SS1 expression relies on the activity of a feed-forward regulatory circuit to activate the master regulator, HilA (50). DMS was implicated as a repressor of HilA because it more potently repressed *hilA* expression than DMSO (34). However, no link was drawn between the ability of *Salmonella* to reduce DMSO and repression of T3SS1 activity. We found that DMSO represses T3SS1 activity even in the absence of all three putative *dmsA* homologs, suggesting that both DMSO and DMS repress T3SS1 activity. One hypothesis for our observation is that there are other reductases in the genome that can convert DMSO to DMS. Therefore, we studied the impact of deletion of *dmsD* in the context of the Δ*dmsA^3^* mutant. The deletion of *dmsD* prevents *dmsA* from binding its cofactor and delivery to the TatABC secretion machinery, thus rendering it non-functional (reviewed in (51)). We also removed the *ynfE* selenate reductase catalytic subunit since *ynfE* is a paralog of *dmsA* and may be able to reduce DMSO (26, 37). Yet, neither of those mutants eliminated the impact of DMSO on T3SS1 repression, suggesting it is unlikely that there is another source of DMS. Therefore, our data are consistent with the hypothesis that both DMSO and DMS impact T3SS1 expression through an unknown mechanism.

We also found that DMSO reduction inappropriately activates expression of the T3SS2. DMSO treatment increased expression of the T3SS2 in low magnesium/low phosphorous medium designed to activate the T3SS2 and in anoxic LB media in which T3SS2 expression is normally deficient (44). These effects were absent in the Δ*dmsA^3^* mutant, suggesting that it is DMS which impacts T3SS2 expression. We were unable to observe an effect of DMS on T3SS2 promoter activity in inducing conditions (data not shown). The boiling point of DMS is 37°C, so it most likely dissipated from liquid culture in the 24-hours used for T3SS2 promoter activation *in vitro*. Although DMSO reduction enhanced T3SS2 expression in media, DMSO reduction did not alter intracellular replication or the proportion of intracellular bacteria expressing a T3SS2 gene, suggesting that the cues influencing T3SS2 expression in the *Salmonella*-containing vacuole are dominant to the impact of DMSO reduction *in vitro*. A mutant which overexpresses the T3SS2 had enhanced replication upon early entry into a macrophage cell line but was attenuated at later time points (52). It is feasible that DMSO could impact intracellular replication at earlier timepoints than we studied here (16 hours). One caveat of our study is that *Salmonella* infection of mammalian cells is conducted under aerobic conditions, which could minimize the impact of DMSO reduction on *S*. Typhimurium pathogenesis *in vitro*. The relevance of DMSO- and DMS-dependent regulation of virulence warrants further mechanistic study.

Our data demonstrate a role for DMSO reduction in colonization of the gut in both murine and bovine hosts. However, we observed differences in the requirement for DMSO reduction between the two animal hosts and between infection models within a host species, underpinning the utility of studying pathogenesis in numerous hosts, and using numerous models in the same host. We observed a necessity for DMSO reduction to support *S*. Typhimurium colonization of the calf gut in the absence of T3SS1-mediated inflammation but no role for DMSO reduction in mice pre-treated with streptomycin, regardless of whether T3SS1-mediated inflammation was present. The finding that DMSO reduction is not needed in mice with antibiotic-induced dysbiosis is consistent with prior work demonstrating no need for *dmsAB* in *E. coli* gut colonization (53). However, we observed a colonization defect for the Δ*dmsA^3^* mutant in the murine host when microbiota were intact. The murine models differ in the time to development of a neutrophilic inflammatory response, the magnitude of the histologic lesions, and the perturbations to the microbiota (31, 32). Therefore, the use of different mouse models allowed us to further interrogate the role of DMSO reduction in prolonged non-inflammatory conditions. We hypothesize that *S*. Typhimurium uses microbiota-derived DMSO to support colonization of the anaerobic gut before higher redox potential electron acceptors (oxygen, nitrate, and tetrathionate) are made available by the robust host inflammatory response. It remains unclear how *Salmonella*-induced enterocolitis alters DMSO availability in the intestine.

*Salmonella* encounters a variety of metabolites produced by the host and the resident gut microbiota during infection. We demonstrated that DMSO reduction is one metabolic process used to support *Salmonella* gut colonization. The three annotated DMSO reductases encoded in the *Salmonella* Typhimurium genome function non-redundantly to support anaerobic growth. Exposure to DMSO and/or DMS also influence the *Salmonella* virulence paradigm, contributing to the complex network of environmental cues regulating virulence. Overall, this work contributes to our understanding of how *Salmonella* gains a foothold in the gut before the onset of severe intestinal inflammation.

## Methods

### Ethics Statement

The Institutional Animal Care and Use Committees of North Carolina State University (protocol numbers 15-047-B (calves) and 17-115-B (mice)) and University of Wisconsin-Madison (protocol number V006255 (mice)) approved all animal experiments. All animal experiments were performed in AALAC-approved animal facilities in accordance with the PHS “Guide for the Care and Use of Laboratory Animals.” All animals used for research were euthanized using AVMA-approved methodology.

### Bacterial strains and growth conditions

All bacterial strains and plasmids are listed in **Table 1**. All bacterial strains are derivatives of ATCC 14028s and construction of mutants was previously described (54). All mutations were moved into a clean genetic background by P22 transduction, antibiotic resistance cassettes removed using FLP recombinase, and mutations confirmed by PCR (55, 56). Unless otherwise specified, bacteria were grown at 37°C in Luria Bertani-Miller (LB) broth with agitation at 225 rpm or on LB agar. Plasmids pMPMA3ΔPlac-null-GFP[LVA], pMPMA3ΔPlac-P*_prgH_*-GFP[LVA], and pMPMA3ΔPlac-P*_ssaG_*-GFP[LVA] were a gift from Olivia Steele-Mortimer (Addgene plasmids # 23338, #23340 and #23343) (35). Bacteria encoding *Salmonella* Typhimurium codon optimized-mCherry under the control of the *trc* promoter were generated by P22 transduction (57). The following antibiotics were used as appropriate: nalidixic acid (50 mg/L), kanamycin (50 mg/L), carbenicillin (100 mg/L), and chloramphenicol (20 mg/L). Dimethyl sulfoxide (DMSO; Fisher Scientific or Sigma) or dimethyl sulfide (DMS; Alfa Aesar) were added to the media at the indicated concentrations.

**Table 1:**
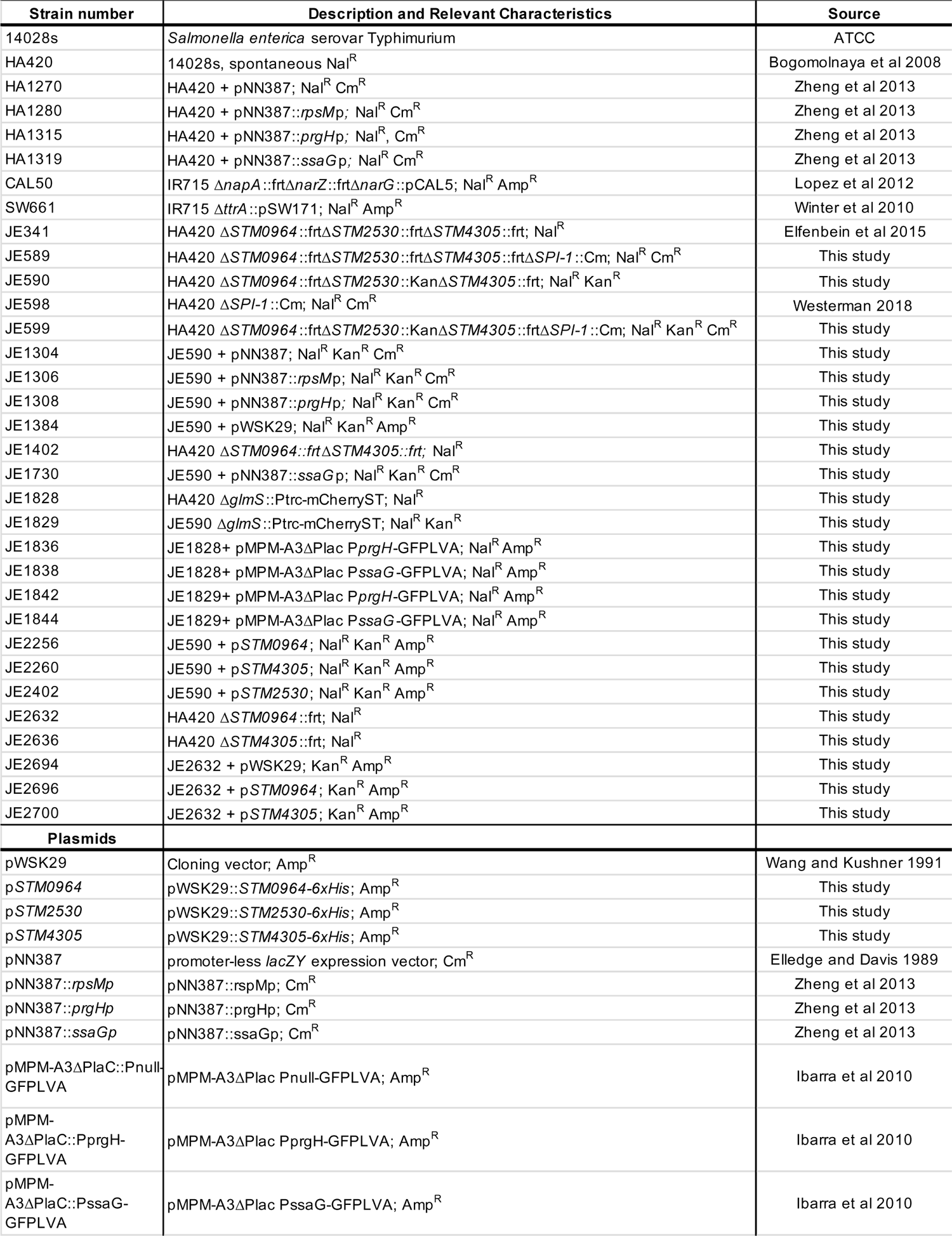
Bacterial strains and plasmids

### Anaerobic and hypoxic growth

For anaerobic growth, overnight cultures were pelleted, supernatants were removed, and cell pellets were transferred into an anaerobic chamber with atmosphere composed of 5% H_2_, 5% CO_2_, and 90% N_2_ (ShelLab, Bactron EZ). Where indicated, overnight cultures were normalized by optical density (600 nm; OD_600_) prior to creating a cell pellet for transfer into the anaerobic chamber. Pellets were re-suspended in LB broth that was pre-reduced in an anaerobic chamber for a minimum of 12 hours and then bacteria were diluted 1:100 into pre-reduced LB broth with the indicated concentrations of DMSO or DMS. Cultures were grown standing at 37°C and aliquots were taken at the indicated times, serially diluted, and plated to determine colony forming units (CFU). Where indicated, the number of generations was determined after 4 hours of growth using the following equation: [log_10_(CFU_4h_) –log_10_(CFU_0h_)]/log_10_(2)]. Each experiment was performed on at least three independent occasions.

For competitive growth at different oxygen tensions, overnight cultures of each bacterial strain were pelleted, resuspended in PBS, and mixed in a 1:1 ratio as determined by OD_600_. Bacterial mixtures were diluted 1:1000 into LB with or without 1% DMSO (v/v) and incubated standing at 37°C in 5% CO_2_ at atmospheric oxygen (normoxic; Eppendorf CellXpert incubator), 1% oxygen (hypoxic; Eppendorf CellXpert incubator), or 0% oxygen (anoxic; ShelLab, Bactron EZ). The ratio of WT to mutant was determined by differential plating of the inoculum and 6-hour cultures. Competitive growth was calculated with the following equation: (WT/mutant)_6h_/(WT/mutant)_0h_. Each assay was performed on 5 independent occasions.

For competitive anaerobic growth in the presence of different electron acceptors, bacteria were mixed in a 1:1 ratio as determined by OD_600_. Mixtures were then pelleted, transferred into the anaerobic chamber, and re-suspended in anaerobic pre-reduced mucin broth. Mucin broth was composed of 0.5% (w/v) hog mucin (Sigma-Aldrich) in no-carbon E media (3.94 g/L KH_2_PO_4_, 5.9 g/L K_2_HPO_4_, 4.68 g/L NaNH_4_HPO_4_*4H_2_O, 2.46 g/L MgSO_4_*7H_2_O) (58). Bacteria were diluted to a starting concentration of ∼10^3^ CFU/mL in mucin broth with DMSO, sodium tetrathionate dihydrate (Sigma-Aldrich), or sodium nitrate (Alfa Aesar), all at a final concentration of 40 mM. Competitive growth was determined after 18 hours. Each assay was performed on at least 3 independent occasions.

### Construction of complementing plasmids

Complementing plasmids each containing an individual *dmsA* homolog, with C-terminal 6x histidine epitope tag, under the control of its native promoter were created using standard cloning protocols. Primer sequences and the corresponding restriction endonucleases are listed in **Table 2**. PCR products were generated by two-step PCR utilizing Q5 polymerase (New England Biolabs) with an annealing temperature of 72°C and a 2-minute extension time. The expected size of PCR products was confirmed by agarose gel electrophoresis and PCR products were purified using QIAquick PCR purification kit (Qiagen). Purified PCR products were sequentially digested with the appropriate restriction endonuclease (New England Biolabs) and re-purified. Gene inserts were cloned into linearized pWSK29 (46) sequentially digested with the appropriate restriction endonucleases and de-phosphorylated with Shrimp Alkaline Phosphatase (New England Biolabs). Inserts were ligated into the linearized plasmid overnight at 16°C using T4 DNA ligase (New England Biolabs) following the manufacturer’s guidelines. Complementing plasmids were transformed into chemically competent DH5α by heat shock at 42°C for 45 seconds. Transformants were selected on LB agar containing carbenicillin and colonies screened for disruption of the multiple cloning site of pWSK29 by blue/white screening with X-gal (5-bromo-4-chloro-3-indolyl-β-D-galactopyranoside; 40 μg/mL). Transformants were purified twice, and the complementing plasmid extracted with QIAprep Spin Miniprep kits (Qiagen). The insert sequence was verified by Sanger sequencing (Functional Biosciences). Each complementing plasmid was transformed into restriction-negative, modification positive *S.* Typhimurium LB5000 by electroporation (59). Transformants were purified twice, plasmids were extracted, and transformed into the appropriate *S*. Typhimurium mutant strains by electroporation. Mutants harboring plasmids were purified twice on LB agar with carbenicillin and stored in glycerol stocks at −80°C.

**Table 2:**
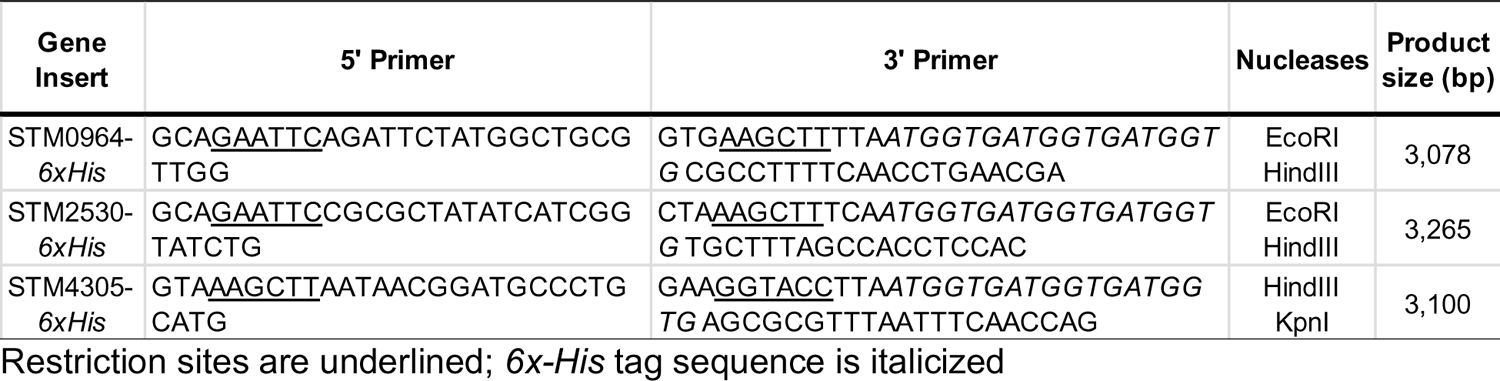
Cloning primer sequences

### Beta-galactosidase assays

Bacteria were transformed with the single-copy number plasmid pNN387 containing a promoter-less *lacZY* or versions of the plasmid with the promoter for *prgH* (type-3 secretion system-1; T3SS1), *ss*a*G* (type-3 secretion system-2; T3SS2) or *rpsM* (ribosomal protein; positive control) driving the expression of *lacZY* (60, 61). For evaluation of T3SS1 gene expression, overnight cultures were diluted 1:100 into LB broth and bacteria were grown at 37°C with agitation for 1.5 hours at which time cultures were treated with 1% (v/v) DMSO or DMS. Bacteria were harvested 1.5 hours after treatment. For measurement of T3SS2 gene expression in inducing conditions, overnight cultures were diluted 1:50 into a low magnesium media [5 mM KCl, 7.5 mM (NH_4_)_2_SO_4_, 0.5 mM K_2_SO_4_, 8 µM MgCl_2_, 337 µM KH_2_PO_4_, 80 mM 2-(N-morpholino)ethanesulfonic acid, 0.3% (v/v) glycerol, 0.1% (v/v) casamino acids, pH 5.8] (43) supplemented with the indicated concentrations of DMSO. Bacteria were harvested after 24 hours incubation standing at 37°C. For measurement of T3SS2 gene expression in non-inducing conditions, overnight cultures were pelleted, transferred into an anaerobic chamber, resuspended in pre-reduced LB and diluted 1:100 into pre-reduced LB with the indicated concentrations of DMSO. Cultures were incubated in an anaerobic chamber at 37°C for 24 hours.

ß-galactosidase activity was quantified using standard methodology (62). Briefly, bacterial cultures were pelleted and resuspended in Z-buffer (60 mM Na_2_HPO_4_, 40 mM NaH_2_PO_4_, 10 mM KCl, 1 mM MgSO_4_, 50 mM ß-mercaptoethanol). Cell density was determined by OD_600_. Cells were permeabilized with chloroform and 0.1% (w/v) SDS followed by addition of 4 mg/mL o-nitrophenyl-ß-D-galactoside (ONPG). Reactions were performed at 28°C and stopped by the addition of 1M Na_2_CO_3_. The absorbances at 420 and 550 nm were determined, and ß-galactosidase activity (Miller units) was calculated using the following equation: 1000 x [OD_420_ – (1.75 x OD_550_)] / [time x volume x OD_600_]. Experiments were performed on at least 3 independent occasions.

### Murine Infection

For the murine colitis model, 8-10-week-old female C57BL/6J mice (Jackson Laboratories, strain# 000664) were used. Mice were treated with 20 mg streptomycin by gavage. Twenty-four hours later, mice were administered ∼10^8^ CFU of an equal mixture of WT and mutant bacteria by gavage and then euthanized 4 days post-infection (32). For the murine oral infection model, 8–10-week-old female CBA/J mice were used (Jackson Laboratories, strain #000656) (30). Mice were administered ∼10^8^ CFU of an equal mixture of WT and mutant bacteria by gavage. Mice were euthanized 8 days post-infection. Following euthanasia, organs were harvested, homogenized, serially diluted, and plated on LB agar with antibiotics as appropriate to quantify CFU. Competitive index was determined by dividing the ratio of the WT to the mutant in the harvested tissues to that of the inoculum.

### Bovine infection

For calf infections, 8 male Holstein, Jersey, or cross-bred calves were obtained from a teaching animal herd at North Carolina State University within 8 hours of birth and housed in individual isolation rooms. Calves were examined by a veterinarian upon acquisition and were treated with ceftiofur (4-6 mg/kg SC q24h for 3-5 days) and/or flunixin meglumine (50 mg/kg IV q24h) as needed based on clinical condition upon arrival. Immediately following acquisition, calves were fed 2-4 L of a commercial colostrum replacer (AgriLabs) and passive transfer of immunity was established 24 hours later by serum total protein measurement. Calves were fed a commercial milk replacer at 10-15% body weight per day with grass hay and water available *ad libitum*. Feces were collected at least twice weekly for selective *Salmonella spp*. culture to ensure all calves remained negative for *Salmonella* fecal shedding.

Between 3 and 6 weeks of age, calves were anesthetized with intravenous propofol and maintained on isoflurane inhalant for ligated ileal loop surgeries as previously described (63). The length of each intestinal segment was measured prior to inoculation with ∼10^9^ CFU containing an equal mixture of the indicated bacterial strains. Infected intestinal segments were harvested 12 hours post-infection and excised to remove luminal fluid. Luminal fluid was weighed and compared with the length of the intestinal segment prior to infection. Loops were separated into fluid, mucus, and tissue fractions. Samples were then homogenized, diluted, and plated to determine CFU. The competitive index was determined by comparing the ratio of two strains in the indicated compartment to the ratio in the inoculum.

### Gentamicin protection assays

HCT116 (human colonic epithelial) cells and J774A.1 (mouse macrophage-like) cells were purchased from ATCC and maintained in McCoy’s 5A media (Corning) containing 10% heat-inactivated fetal calf serum (FCS, Invitrogen) or Dulbecco’s Modified Eagle’s medium (Corning) containing 10% heat-inactivated FCS, respectively. Cells were used within 15 passages of receipt from ATCC. Cells were seeded in 24-well plates at 1×10^5^ cells/well (HCT116) 44-48 hours prior to infection or 2.5×10^5^ cells/well (J774A.1) 24 hours prior to infection. Bacterial subcultures were prepared from overnight cultures (16-18 hours growth in LB-Miller with shaking at 37°C) by back-diluting 1:33 and shaking in LB-Miller broth in Erlenmeyer flasks for 3 hours at 37°C. For DMSO treatment, 1% (v/v) DMSO was added to subcultures after 1.5 hours of growth and added to tissue culture media to 1% (v/v) after the initial 10 minutes infection period and maintained for the duration of the infection. Infection of HCT116 cells and J774A.1 cells was as described previously (64). At 1 hour or 16 hours post-infection, infected monolayers were washed once in phosphate-buffered saline (PBS), solubilized in 1 ml 0.2% sodium deoxycholate, and serial dilutions were plated on LB agar for CFU enumeration.

### Fluorescence microscopy

HCT116 cells and J774A.1 cells were seeded on 12 mm acid-washed glass coverslips in 24-well plates as described above. At the required timepoints, infected monolayers were washed once with PBS, fixed with 2.5% (w/v) paraformaldehyde prepared in PBS for 10 minutes at 37°C. DNA was stained with Hoechst 33342 (Invitrogen) and coverslips were mounted onto glass slides using Mowiol. The proportion of GFP-positive bacteria was scored by fluorescence microscopy by an investigator blinded as to sample identification.

### Statistical analyses

Data were analyzed using GraphPad Prism 9 v9.2. Competitive indices were log transformed prior to analysis. Student’s t-test or two-way ANOVA were used where indicated. A repeated-measures two-way ANOVA was used to analyze calf infection data. Corrections for multiple comparisons were made as indicated. Significance was set at P<0.05 for all analyses.

## Acknowledgements

This work was supported, in part, by start-up funds from the University of Wisconsin-Madison and Food Research Institute to JRE. JRE was supported, in part, by NIH K08AI108794. JRE and LAK were supported, in part, by USDA-NIFA 2018-67017-27632. EC acknowledges support from an American Society for Microbiology Undergraduate Research Fellowship, a Foundation for Food and Agricultural Research Veterinary Fellowship, and a University of Wisconsin-Madison Science and Medicine Graduate Research Fellowship.

